# *Melanoides tuberculata* in East Africa: Investigating the role of Biotic and Abiotic factors in its presence or absence

**DOI:** 10.64898/2026.07.28.741188

**Authors:** Victor O. Magero, Sammy Kisara, Christopher M. Wade

## Abstract

**Background:** *Melanoides tuberculata* is one of the most highly invasive snails in the world with strong colonizing capabilities. Its competitive advantage over other freshwater snails has led to its use to biologically control snails that act as intermediate hosts of *Schistosoma mansoni*, averting the spread of schistosomiasis (a Neglected Tropical Disease infecting approximately 40 million people in East Africa). In East Africa, few studies have been conducted to better understand the biotic and abiotic factors that promote the survival and dominance of *M. tuberculata* snails. This study purposed to investigate the role of temperature, soil pH, water-depth, water-velocity, soil type, vegetation cover and the presence or absence of other snails on the presence or abscence of *M. tuberculata* snails from locations across East Africa.

**Results:** Water depth (p=0.001) was found to be the most statistically significant factor affecting the presence of *M. tuberculata* snails at the East African sites; the average water-depth where *M. tuberculata* snails were found was 20.7cm. The presence or absence of other snails was also found to have a statistically significant role on the presence of *M. tuberculata* snails (p=0.005), with *M. tuberculata* being rarely found alongside other snails. Water temperature also had a statistically significant role on the presence of *M. tuberculata* snails (p=0.014); the average water temperature where *M. tuberculata* snails ere found was 20.27°C.

Vegetation cover (p=0.236), pH (p=0.434), water-velocity (p=0.736) and soil type (p=0.356) potentially had a role on the presence or absence of East African *M. tuberculata* snails but their effect was not found to be statistically significant. All 7 locations where *M. tuberculata* snails were found had vegetation cover, an average pH of 8.07, an average water-velocity of 0.49m/s and silt and sandy soil as the soil type.

**Conclusion:** In conclusion, we found that water depth, water temperature and the presence or absence of other snails had a significant role on the presence of *M. tuberculata* snails in East Africa. Our study provides detailed information on the role of biotic and abiotic factors on the presence of *M. tuberculata* snails in several locations of East Africa.

## 1.0 INTRODUCTION

*Melanoides tuberculata* is a freshwater caenogastropod snail that is considered to be one of the most highly invasive snail species in the world (Pinto *et al*., 2023). The snail reproduces sexually and via parthogenesis (making it possible for one individual to produce a population of high density within a short period of time), has fast growth, is an omnivore and has a wide tolerance to different environmental stressors (Durán-Rodríguez *et al*., 2024). *M. tuberculata* snails have colonizing behaviour (they often turn new areas into their own territories) and tend to prefer regions that have high human activities (Durán-Rodríguez *et al*., 2024). Several studies affirm that *M. tuberculata* has the ability to displace native gastropods (Murray, 1971; Jacobson, 1975; Hershler, 1998). The snails live for relatively long periods; 12-15 months in their natural state, with some individuals being reported to live as long as 2-3 years (Kadri, 1974; Leveque, 1973; Abell & Nyamweru, 1988). *M. tuberculata* is very advantageous in improving soil air quality and consequently soil health because as the snails move in the soil eating organic waste and litter, they aerate the soil thus promoting root growth (Abell & Nyamweru, 1988).

*M. tuberculata* is native to Sub-Saharan Africa and South Asia (Abell & Nyamweru, 1988) but multiple introductions, mainly trade in aquarium plants and animals has played an important role in the introduction of the snail to places as far as Europe, America and Oceania (Pinto *et al*., 2023). Brown (1980) asserts that *M. tuberculata* is the most commonly collected caenogastropod species, in Kenya and that it is frequently found in the same habitats that support the presence of *Biomphalaria pfeifferi* (an important snail intermediate host of *Schistosoma mansoni*) and *Lymnaea natalensis* (an important snail intermediate host of *Fasciola gigantica*).

*M. tuberculata*’s adult sizes in native regions range between 20-40 mm but sizes of up to 80mm have been recorded. Abiotic factors that influence the presence of the snails include temperature, chemistry of water (water hardness, salinity, pH) and conditions of the microhabitat (it has been shown to be absent in fast flowing waters) (Hiba & Zainab, 2022).

*M. tuberculata* has been demonstrated to show promise as a biological tool in the control of snail intermediate hosts, in the fight against schistosomiasis (Pointier & McCullough, 1989). *M. turberculata* has been introduced to the Caribbean and Neotropics in an effort to control freshwater snails, notably *Biomphalaria* snails that serve as intermediate hosts of *Schistosoma* parasites (Pointier & McCullough, 1989; Butlers *et al*., 1980; Pointier *et al*., 1989). A 10 year study reported that *M. tuberculata* was able to eradicate *Biomphalaria glabrata* and *Biomphalaria straminea* from two lakes in Brazil eight years after the introduction of *M. tuberculata* (Guimaraes *et al*., 2001).

*M. tuberculata* is highly invasive and can outcompete snail intermediate hosts that transmit schistosomiasis (Pinto *et al*., 2023). Other than serving as a competitor snail, *M. tuberculata* has been shown to be an intermediate snail host for over 40 trematode species including a fluke that infects humans, fish and the fountain darter (*Centrocestus formosanus*,); eye flukes that infect birds (*Philophthalmus* species); intestinal flukes that infect snails, fish, birds, humans and other mammals(*Haplorchis taichui* and *Haplorchis pumilio*); a flatworm that infects humans and animals (*Echinochasmus milvi*) and several blood flukes (Pinto & Melo, 2011).

In East Africa, a region shown to harbour schistosomiasis hotspots, few studies have been conducted to understand the biotic and abiotic factors that promote the survival and dominance of *M. tuberculata*. This study purposed to investigate biotic and abiotic factors that support the presence or absence of *M. turberculata* in East Africa.

## 2.0 METHODS AND METHODOLOGY

### 2.1. Malacological surveys

Extensive malacologcal surveys were carried out in East Africa between 2018 – 2020, with 172 sites surveyed across Kenya, Uganda and Tanzania for the purpose of finding *M. tuberculata* snails (Figure 1; Supplementary Table 1). The surveys involved spending approximately 20 minutes at every given site looking for live snails and searching for any cues for the availability of snails such as dead shells. All snails present at a site were collected and identification keys developed by Brown (1994) were used to identify *M. tuberculata* and other snails based on their morphological characteristics. The snails were collected using a wiremesh, handpicked with the aid of forceps and transferred into falcon tubes containing absolute ethanol to preserve the snails.

**Figure 1.**
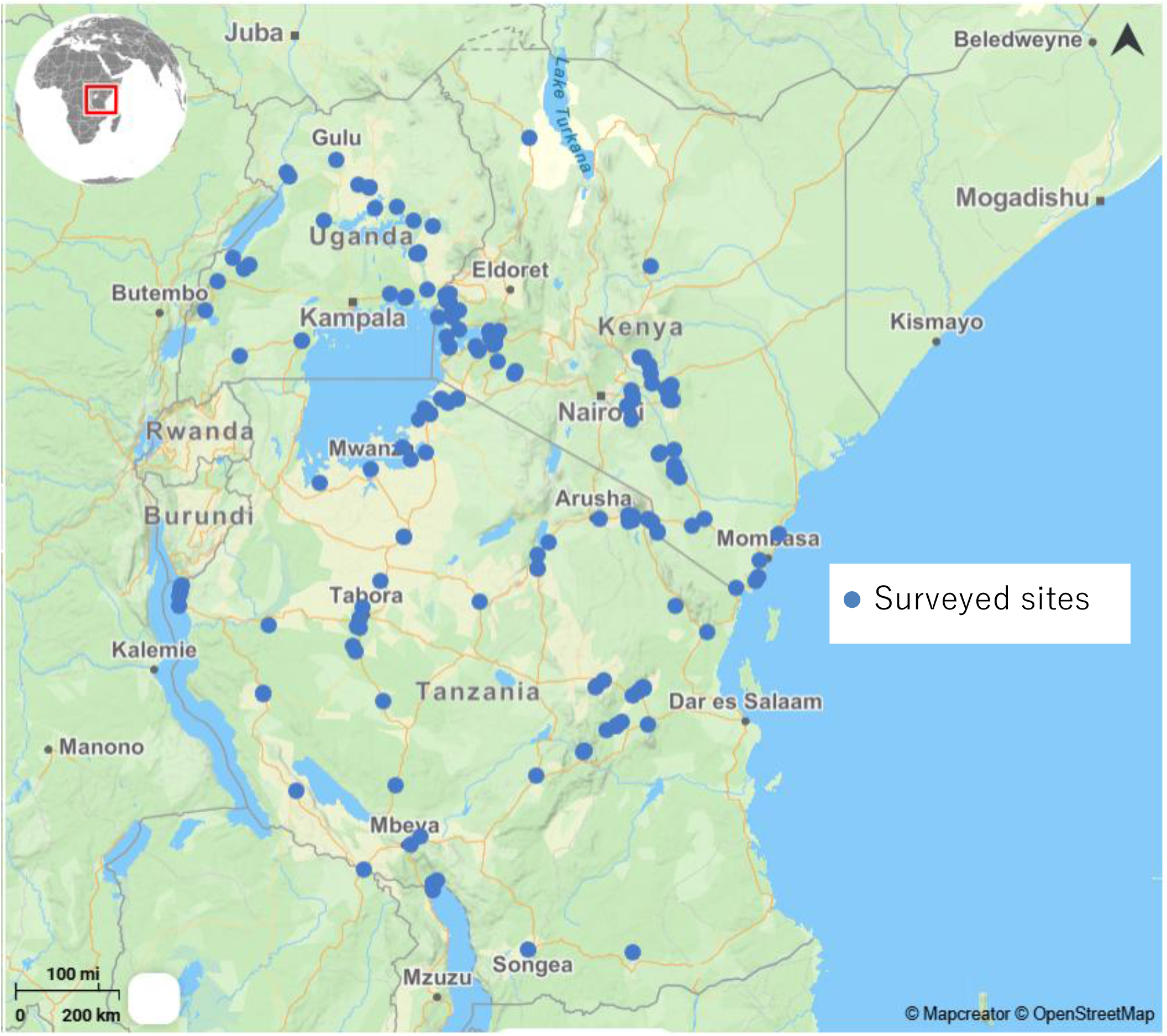
Map of East Africa showing sites visited for Malacological survey purposes, in search of *M. tuberculata* snails. The map was created using a basemap from OpenStreetMap

### 2.2 Measurement of biotic and abiotic factors

At each of the 172 sites, physical variables of habitat and physicochemical parameters were measured. Water temperature and soil pH were recorded using a portable water metre (Hanna Instruments, Møllevænget, Sweden). Water depth was recorded using a 1 metre ruler and water velocity was recorded using a flowmeter, which turns with the flow of a stream or a river (Ultrasonic Flow Meter, Doyi, China). The soil type for each site was noted and recorded was the vegetation cover for each site. Finally the presence or absence of other snails at each site was recorded.

Statistical analysis was conducted to ascertain the relationship that exists between physical variables of habitat and physicochemical parameters with the presence or absence of *M. tuberculata* snails at the 172 East African sites. Pearson correlation coefficient and Spearman’s rank correlation coefficient in SPSS (Statistical Package for Social Sciences) version 31 was used to test the relationship between physicochemical parameters (biotic and abiotic factors) and the presence of *M. tuberculata* snails.

## 3.0 RESULTS

### 3.1 Locations where *M. tuberculata* snails were found

*M. tuberculata* snails were found at just 7 sites of the total 172 sites surveyed in East Africa: Kosena (0.09, 34.28), Mutendea (-1.33, 37.98), Macheche (0.39, 33.48), Ntoroko (1.05, 30.54), Kikavu (-3.44, 37.29), Wang-Lei (2.26, 32.88) and Kabere (0.41, 33.50) (Table 1, Supplementary Table 1, Figure 2, Figure 3).

**Table 1:** Sites where *M. tuberculata* snails were found, with their respective biotic and abiotic factors.

| Name of site | Geographic Coordinates | Type of habitat | Water Temp (° C) | Soil pH | Water depth (cm) | Water velocity (m/s) | Soil Type | Vegetation (Yes/No) | Presence or absence of other snails |
| --- | --- | --- | --- | --- | --- | --- | --- | --- | --- |
| Kosena (Kenya) | (0.09, 34.28) | Dam | 15.3 | 8.1 | 19.2 | 0.6 | Sandy soil | No | No |
| Mutendea (Kenya) | (-1.33, 37.98) | Stream | 29.1 | 7.3 | 35.2 | 0.5 | Sandy soil | Yes | No |
| Macheche (Uganda) | (0.39, 33.48) | Dam | 26.2 | 8.1 | 16.3 | 0.8 | Sandy soil | Yes | No |
| Ntoroko (Uganda) | (1.05, 30.54) | River | 20.8 | 8.4 | 11.2 | 0.6 | Silt soil | Yes | Yes |
| Kikavu (Tanzania) | (-3.44, 37.29) | Irrigation scheme | 16.2 | 8.2 | 12.4 | 0.2 | Sandy soil | Yes | No |
| Wang Lei (Uganda) | (2.26, 32.88) | Stream | 15.2 | 7.2 | 18.6 | 0.3 | Silt soil | Yes | No |
| Kabere (Uganda) | (0.41, 33.50) | Stream | 19.1 | 9.2 | 27.6 | 0.4 | Silt soil | Yes | No |

**Figure 2.**
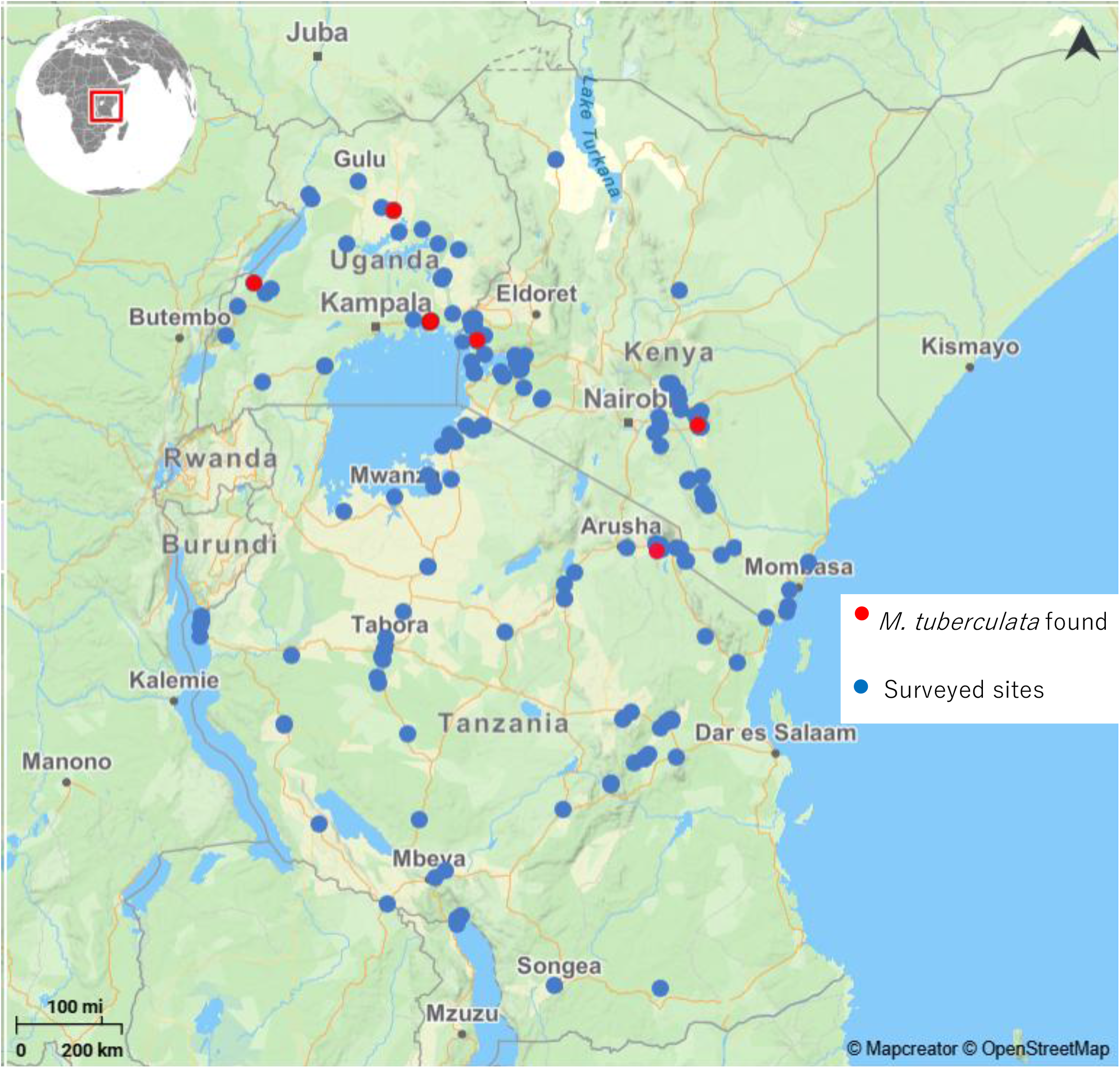
Map of East Africa showing surveyed sites and locations where *M. tuberculata* snails were found. The map was created using a basemap from OpenStreetMap.

**Figure 3.**
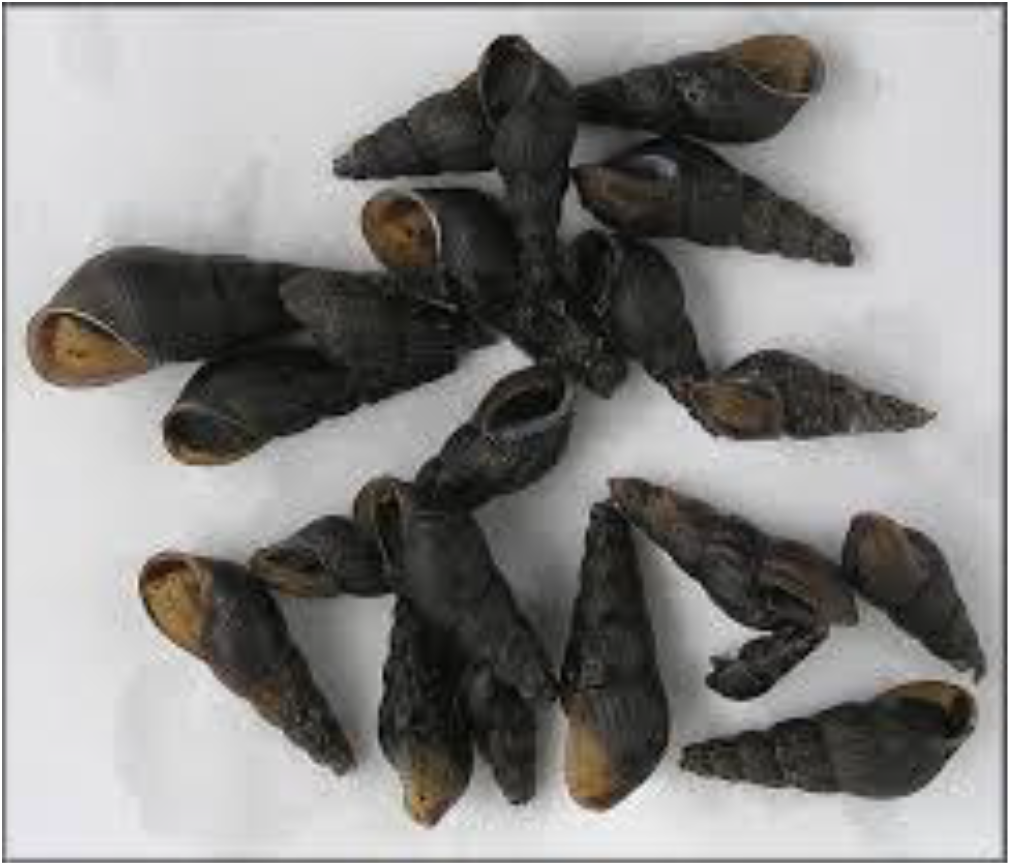
Image of *M. tuberculata* snails collected from Kikavu irrigation scheme (Tanzania)

### 3.2 Abiotic and biotic factors that affect the presence or absence of *M. tuberculata* snails

The biotic and abiotic factors whose role in the presence or absence of *M. tuberculata* snails were investigated were water temperature, pH, water-depth, water-velocity, soil type, vegetation cover and the presence and absence of other snails. Water depth (p=0.001) was found to be the most statistically significant factor that affects the presence or absence of *M. tuberculata* in East Africa. The average water-depth for the locations where *M. tuberculata* snails were found was 20.7cm (range 11.2 – 27.6cm) when compared to an average water depth of 31.0cm (range 12.5 – 53.2cm) at the sites where *M. tuberculata* snails were not found. Likewise, water temperature had a statistically significant role in the presence of *M. tuberculata* snails (p=0.014). The average water temperature for the locations where the *M. tuberculata* snails were found was 20.27°C (range 15.1 – 29.5°C) when compared to an average water temperature of 22.5°C (range 13.2 – 32.3°C) for the locations where *M. tuberculata* snails were not found.

The presence or absence of other snails was also found to have a statistically significant role in the presence of *M. tuberculata* snails at a site (p=0.005). Of the 7 sites where *M. tuberculata* snails were found, no other snail species were present at 6 of the 7 sites. *M. tuberculata* snails were found alongside other snails at a single site (Ntoroko) where only 5 *Biomphalaria* snails were found, and where hundreds of *M. tuberculata* snails were found amidst hundreds of dead *Biomphalaria* shells.

All locations where *M. tuberculata* snails were found had vegetation cover, a silt and sandy soil as the soil type, an average soil pH of 8.07 (range 7.2 – 9.2), and an average water velocity of 0.49m/s (range 0.2 – 0.8m/s). However, vegetation cover (p=0.236), soil type (p=0.356), PH (p=0.434) and water-velocity (p=0.736) were not found to have a statistically significant effect on the presence or absence of *M. tuberculata* snails.

## 4.0 DISCUSSION

This study established that biotic and abiotic factors potentially have an influence on the presence or absence of *M. tuberculata* snails in East African locations. Water depth was found to be the most statistically significant abiotic factor that affects the presence of *M. tuberculata* snails (p=0.001), across the 172 sites sampled. The average water-depth for the locations where *M. tuberculata* snails were found was 20.7cm. In the USA, *M. tuberculata* snails have been found to be abundant in shallow sites where water depth is less than 30 cm; sites known to have an abundance of food (Anderson, 2004). Drastic water fluctuations are likely to lead to a decline in the presence of *M. tuberculata* snails, given that most of them survive in shallow waters (Hunt & Jones, 1972). A study of *M. tuberculata* in Brazil found high snail densities in water depths between 20cm and 30cm (Oliveira & Oliveira, 2019). Schmitt *et al*. (2024) states that *M. tuberculata* snails from Oman prefer aquatic environments with water depth of up to 25 cm.

Water temperature was also demonstrated to have a statistically significant role on the presence of *M. tuberculata* snails (p=0.014). The average water temperature for the locations where the *M. tuberculata* snails were found was 20.27°C. *M. tuberculata* snails have been shown to have a preference for warm waters of a temperature range between 18 - 31 C^°^(Van-Bocxlaer *et al*., 2025). Warmer temperatures have been shown to increase metabolic activities and growth rates of the snails, promoting their invasive capability in South-Africa (De Kock & Wolmarans, 2009), in New-Zealand (Duggan, 2002) and in Argentina (Seuffert *et al*., 2023). Van-Bocxlaer *et al*. (2025) reported an increase in size and growth of juvenile *M. tuberculata* snails, with increase in temperature at Lake Malawi. A study by Okumura & Rocha (2020) on *M. tuberculata* kept in the laboratory showed a narrow range of temperature tolerance (between 16°C to 37°C). Bede (1992), observed an increase in the growth and reproduction of *M. tuberculata* snails from Brazil at higher temperature ranges between 29°C to 33°C. When temperatures exceed 35°C, *M. tuberculata* snails in Utah (USA) have been shown to become nocturnal, spending their time in mud (Rader *et al*., 2003). When temperatures in an experimental setting were increased over 32°C, mortality rates among the juvenile *M. tuberculata* snails increased (Van-Bocxlaer *et al*., 2025). Water temperature has also been shown to play a role on the amount of resources that *M. tuberculata* snails use to invest in their immune system (Van-Bocxlaer *et al*., 2025). At cold temperatures, the snails use high proportions of their resources to boost their immune system whereas in warm environments, the snails did not heavily invest their resources in immune systems, thus the said resources are alternatively used for growth, reproduction and dispersal purposes (Van-Bocxlaer *et al*., 2025).

Soil type has a potential role on the presence of *M. tuberculata* snails, though the effect of soil type on the presence of *M. tuberculata* snails in our East African study was not found to be statistically significant (p=0.356). In our study, *M. tuberculata* snails were found at sites with silt and sandy soils and were not found at sites with muddy, rocky or gravel soils. *M. tuberculata* snails in French West Indies have been associated with different types of soil substrates such as sandy soil, clay soil, mud, gravel and rocky soil (Vogler *et al*., 2012). A study by Schmitt *et al*. (2024) in Oman suggests that *M. tuberculata* snails prefer moist soil with a moderate amount of humus.

Soil pH also has a potential role on the presence of *M. tuberculata* snails, but again the relationship between soil pH and the presence of *M. tuberculata* snail was not found to be statistically significant (p=0.434) in our study. In previous studies, *M. tuberculata* snails have been shown to have a pH preference of 7.0 -7.5 and lower pH values have been found to reduce the densities of the snails (Vogler *et al*., 2012; Dudgeon, 1986; Liyanagedara *et al*., 2023). pH of less than 5 has been reported to be deleterious to the growth and survival of *M. tuberculata* snails in Martinique (Vogler *et al*., 2012). pH levels less than 5 have been reported to make it difficult for snails to uptake calcium that is necessary for shell construction (the snails requiring high energy expenditure, in the process). Extremely low pH levels (as low as 4.0 or 2.0) have been reported to be lethal to the growth and survival of embryos and juvenile snails in Martinique and Hong-Kong (Vogler *et al*., 2012; Dudgeon, 1986). A study by Al-Baghdadi *et al*. (2023) in Iraq affirmed that high pH values (alkaline waters) are ideal for the development of *M. tuberculata* shells. pH has also been established to be one of the most important abiotic factors in the presence or absence of freshwater snails in Cuba (Gutiérrez *et al*., 1997). Calcium levels have been reported to contribute towards pH levels in aquatic ecosystems, with high calcium concentrations playing a role in increasing pH levels in Connecticut (USA) (Jokinen, 1983).

Water velocity, potentially also has a role on the presence of *M. tuberculata* snails, but again the relationship between water velocity and snail presence was not statistically significant (p=0.736) in our study. The majority of sites visited in our study had slow flowing waters including the sites where *M. tuberculata* snails were found. In Martinique, water velocities higher than 1.1 m/s were found to discourage the presence of *M. tuberculata* snails (Dussart & Pointier, 1999). According to Vogler *et al*. (2012), *M. tuberculata* snails, in West Indies have been found to reach high densities of abundance in gently flowing waters. Vogler *et al*. (2012) asserts that *M. tuberculata* snails have been found in rivers and streams with different flow rates, but completely absent in water bodies with steep, fast-flowing waters.

Biotic factors are also potentially important in the growth, development and presence of *M. tuberculata* snails. The presence/absence of other snails in locations where *M. tuberculata* snails were present and the presence of vegetation cover are the two biotic factors that were investigated in this study. The presence/absence of other snails was found to have a statistically significant role on the presence or absence of *M. tuberculata* snails in the East African locations (p=0.005). *M. tuberculata* was found alongside other snail species at only 1 of the 7 sites where *M. tuberculata* snails were found (at Ntoroko, Uganda). However, only 5 *Biomphalaria* snails were found at Ntoroko, amidst hundreds of *M. tuberculata* snails and hundreds of dead *Biomphalaria* shells. The presence of hundreds of dead *Biomphalaria* shells at Ntoroko could be interpreted as *M. tuberculata*’s colonizing ability and capability to displace other snails from any given habitat. A study by Rader *et al*. (2003), revealed *M. tuberculata*’s competitive advantage in spring ecosystems of the Bonneville Basin, Utah (USA), with the snail becoming one of the most abundant snail species 5 to 8 years after colonizing 2 springheads in a large spring complex. In Burkina Faso, *M. tuberculata* has also been documented to have a strong competitive advantage over other snails, having colonised 10 out of the 12 surveyed sites (Sankara *et al*., 2026). The competitive advantage that *M. tuberculata* has over other snails has been attributed to its parthenogenetic reproduction, being tolerant to high salinity levels, fast-growth, tolerance to disturbed habitats (including those affected by anthropogenic activities) and being an omnivore feeder (Durán-Rodríguez *et al*., 2024; (Pinto *et al*., 2023).

Vegetation cover also plays a potential role on the presence of *M. tuberculata* snails in East Africa, though the relationship between vegetation cover and the presence and absence of *M. tuberculata* snails was not found to be statistically significant (p=0.236) in our study. Vegetation cover was found at all 7 sites where *M. tuberculata* snails were found in our study, but vegetation cover was also found at many sites where *M. tuberculata* snails were not found. Vegetation cover has been shown to be important in providing shedding and foliar material for the establishment of *M. tuberculata* snails in new habitats of Mexico (Durán-Rodríguez *et al*., 2024) and Brazil (Barros *et al*., 2020). Vegetation has previously been shown to be important in influencing the presence of freshwater snails in Kenya as the snails use the vegetation for cover (shedding), egg laying, source of food and for attachment purposes (Dida *et al*., 2014).

Anthropogenic alterations have been found to promote the invasion and dominance of *M. tuberculata* snails in new habitats of Brazil (Barros *et al*., 2020). A study by Weir & Salice (2011) on *M. tuberculata* snails fromTexas (USA) suggested that *M. tuberculata* had a strong advantage over *B. glabrata* in disturbed and polluted environments. Water salinity has also been established to play an important role in the presence or absence of *M. tuberculata* in the Chalbi Basin of Kenya (Abell & Nyamweru, 1988), in the Ceará State of Brazil (Barros *et al*., 2020) and in French West Indies (Vogler *et al*., 2012). We did not explore the role of anthropogenic alterations or salinity on the presence or absence of *M. tuberculata* snails in our study.

## 5.0 CONCLUSION

In conclusion, we found that water depth, water temperature and the presence or absence of other snails had a significant role on the presence of *M. tuberculata* snails in East Africa. Our study provides detailed information on the role of biotic and abiotic factors on the presence of *M. tuberculata* snails in several locations of East Africa.

## List of abbreviations

pH: potential of hydrogen; a measure of acidity and alkalinity
cm: centimetre
mm: millimetre
SPSS: Statistical Package for Social Sciences
m/s: metres per second
^°^ C: degree Celsius

## DECLARATIONS

## Acknowledgements

We would like to acknowledge Liaque Latif and Sara Goodacre for their input in the success of this study.

## Authors’ Contributions

Victor O. Magero was involved in conceptualisation, methodology, investigation, data curation, formal analysis and writing the original draft.

Sammy Kisara was involved in methodology, investigation and data curation.

Christopher M. Wade was involved in conceptualisation, methodology, project administration, validation, formal analysis and review and editing the manuscript.

## Ethical Approval

Ethical approval not required

## Consent to Participate

Consent to participate, not required

## Consent to Publish

Consent to publish not required

## Data availability statement

The data that support the findings of this study are available in the attached Supplementary file.

