## Supplemental Table 1 for "*Melanoides tuberculata* in East Africa: Investigating the role of Biotic and Abiotic factors in its presence or absence"

**Supplementary Table 1:** Geographic coordinates; biotic and abiotic factors data.

| **Name of site** | **Coordinates** | **Type of habitat** | **Water Temp (°C)** | **pH** | **Water depth (cm)** | **Water velocity (cm/s)** | **Soil Type** | **Vegetation**  **(Y/N)** | **Presence/absence of other snails (Y/N)** | ***M. tuberculatafound* (Y/N)** |
| --- | --- | --- | --- | --- | --- | --- | --- | --- | --- | --- |
| Mbondoni (Kenya) | (-0.97325, 38.005) | Dam | 31.2 | 7.3 | 12.5 | 0.9 | Sandy soil | N | N | N |
| Migwani (Kenya) | (-1.09305556, 38.02111) | Dam | 29.1 | 6.2 | 15.4 | 0.1 | Sandy soil | N | N | N |
| Matingani (Kenya) | (-1.164, 38.005) | Spring | 23.2 | 5.2 | 13.7 | 0.3 | Clay soil | Y | Y | N |
| Kangonde (Kenya) | (-1.07941667,  37.69166667) | Dam | 31.2 | 8.3 | 26.4 | 0.7 | Sandy soil | N | N | N |
| Mutendea (Kenya) | (-1.33180556,  37.98166667) | Stream | 29.1 | 7.3 | 35.2 | 0.5 | Sandy soil | Y | N | Y |
| Kiangangi (Kenya) | (-0.593833,  37.341972) | Irrigation scheme | 24.5 | 8.2 | 17.5 | 0.5 | Silt soil | Y | Y | N |
| Kitui (Kenya) | (-1.35616667,  38.00611111) | Stream | 20.6 | 6.4 | 16.3 | 0.4 | Silt soil | Y | Y | N |
| Kalundu (Kenya) | (-1.35616667,  38.00611111) | Stream | 20.4 | 7.3 | 23.1 | 0.5 | Sandy soil | Y | Y | N |
| Kalundu (Kenya) | (-1.36555556,  38.00277778) | Stream | 20.1 | 8.2 | 19.4 | 0.1 | Sandy soil | Y | Y | N |
| Ikindu  (Kenya) | (-1.36733333,  38.03916667) | Stream | 31.2 | 9.2 | 28.2 | 0.7 | Sandy soil | Y | N | N |
| Nzeu (Kenya) | (-1.37061111, 38.02416667) | Stream | 29.1 | 7.2 | 12.1 | 0.2 | Sandy soil | N | N | N |
| Kwase (Kenya) | (-1.29883333,  37.35722222) | Stream | 19.3 | 6.8 | 19.3 | 0.5 | Silt soil | Y | Y | N |
| Mutanga (Kenya) | (-1.35758333,  37.35472222) | Stream | 25.7 | 8.2 | 25.1 | 0.3 | Silt soil | Y | Y | N |
| Chumbe (Kenya) | (-2.26511111, 37.81027778) | Stream | 32.1 | 7.9 | 19.3 | 0.6 | Sandy soil | N | N | N |
| Ikoyo (Kenya) | (-2.26613889, 37.81) | Dam | 31.2 | 8.4 | 25.1 | 0.3 | Sandy soil | N | N | N |
| Thange (Kenya) | (-2.47025,  38.06583333) | River | 14.1 | 6.8 | 29.6 | 0.2 | Sandy soil | Y | N | N |
| Thange (Kenya) | (-2.50158333,  38.02277778) | River | 15.6 | 7.3 | 22.4 | 0.5 | Sandy soil | Y | N | N |
| Thange (Kenya) | (-2.47536111,  38.09527778) | River | 15.2 | 8.3 | 21.3 | 0.6 | Sandy soil | Y | N | N |
| Kambu (Kenya) | (-2.536591, 38.118119) | River | 31.5 | 7.4 | 22.8 | 0.6 | Sandy soil | Y | N | N |
| Kambu (Kenya) | (-2.498523, 38.053190) | River | 30.9 | 9.3 | 29.2 | 0.4 | Sandy soil | Y | N | N |
| Loilopon (Kenya) | (-2.930847, 37.476999) | Stream | 13.2 | 8.2 | 32.4 | 0.6 | Silt soil | N | N | N |
| Kambu (Kenya) | (-2.56630556,  38.11944444) | River | 20.9 | 9.3 | 33.6 | 0.3 | Sandy soil | Y | N | N |
| Itua (Kenya) | (-0.62888889,  37.54) | River | 25.4 | 8.3 | 28.3 | 0.2 | Silt soil | Y | Y | N |
| Thiba (Kenya) | (-0.77269444,  37.63888889) | River | 15.3 | 7.2 | 35.2 | 0.4 | Sandy soil | Y | N | N |
| Tulimiumbu (Kenya) | (-0.91133333,  37.65833333) | Stream | 21.3 | 6.8 | 23.1 | 0.5 | Sandy soil | Y | Y | N |
| Kitunene (Kenya) | (-0.92533333,  37.67583333) | Dam | 15.2 | 9.2 | 36.3 | 0.1 | Sandy soil | N | N | N |
| Musilili (Kenya) | (-1.45023611,  37.2575) | Stream | 24.6 | 8.3 | 25.2 | 0.5 | Silt soil | Y | Y | N |
| Kakulutuine (Kenya) | (-1.20472222,  37.33055556) | Stream | 25.3 | 7.6 | 21.7 | 0.3 | Silt soil | Y | Y | N |
| Mukou (Kenya) | (-1.68488889,  37.34472222) | Stream | 25.9 | 9.2 | 19.2 | 0.5 | Silt soil | Y | Y | N |
| Mtito (Kenya) | (-2.68713889, 38.16) | River | 21.2 | 8.1 | 28.2 | 0.2 | Sandy soil | Y | N | N |
| Kahoma (Kenya) | (-3.42088889, 37.68305556) | Stream | 15.4 | 8.2 | 27.2 | 0.6 | Silt soil | Y | N | N |
| Voi (Kenya) | (-3.39063889,  38.57861111) | River | 13.2 | 6.4 | 33.2 | 0.3 | Sandy soil | N | N | N |
| Jipe A (Kenya) | (-3.59861111,  37.77444444) | Stream | 31.5 | 7.3 | 33.4 | 0.6 | Sandy soil | Y | N | N |
| Jipe B (Kenya) | (-3.61477778,  37.77611111) | Stream | 32.1 | 8.2 | 28.9 | 0.4 | Sandy soil | Y | N | N |
| Jombo (Kenya) | (-3.50175, 38.37777778) | River | 14.2 | 7.1 | 25.2 | 0.2 | Sandy soil | N | N | N |
| Kasabong (Kenya) | (-0.15608333, 34.40583333) | Stream | 29.3 | 8.3 | 32.1 | 0.2 | Silt soil | N | N | N |
| Abitha (Kenya) | (-0.16761111,  34.40583333) | Stream | 29.7 | 9.1 | 15.2 | 0.7 | Silt soil | Y | N | N |
| Gera (Kenya) | (-0.46413889,  34.22916667) | River | 31.2 | 8.2 | 28.2 | 0.6 | Silt soil | Y | N | N |
| Nyabanda (Kenya) | (-0.08961111,  34.28111111) | Dam | 15.3 | 9.1 | 25.4 | 0.7 | Sandy soil | Y | N | N |
| Manyonge (Kenya) | (-0.06366667,  34.29166667) | Stream | 14.2 | 7.8 | 36.2 | 0.7 | Silt soil | Y | N | N |
| Nyawita (Kenya) | (-0.43036111,  34.15472222) | Stream | 13.5 | 9.2 | 19.2 | 0.6 | Silt soil | Y | N | N |
| Kosena (Kenya) | (0.09338889, 34.28) | Dam | 15.3 | 8.1 | 19.2 | 0.6 | Sandy soil | N | N | Y |
| Ambururu (Kenya) | (0.15163889,  34.27916667) | Stream | 23.5 | 7.5 | 30.2 | 0.4 | Silt soil | Y | Y | N |
| Mbita (Kenya) | (-0.42180556,  34.20611111) | River | 29.3 | 6.3 | 39.6 | 0.7 | Sandy soil | Y | N | N |
| Thogoye (Kenya) | (-0.06091667,  34.04305556) | River | 31.2 | 7.2 | 30.2 | 0.9 | Sandy soil | Y | N | N |
| Gera River (Kenya) | (-0.44213889,  34.22666667) | River | 30.4 | 8.3 | 29.1 | 0.4 | Sandy soil | Y | N | N |
| Luanda (Kenya) | (-0.46413889,  34.22916667) | Stream | 29.2 | 7.5 | 29.2 | 0.4 | Silt soil | Y | N | N |
| Kulo Kudongo (Kenya) | (0.27072222,  34.19083333) | Stream | 31.3 | 6.9 | 32.1 | 0.3 | Silt soil | Y | N | N |
| Wathi (Kenya) | (0.30513889,  34.23472222) | River | 30.5 | 7.2 | 33.3 | 0.7 | Sandy soil | Y | N | N |
| Rakite (Kenya) | (0.32575,  34.19472222) | Stream | 21.2 | 8.2 | 23.1 | 0.4 | Silt soil | Y | Y | N |
| Matodo (Kenya) | (0.30847222,  34.22972222) | Stream | 31.3 | 7.3 | 32.1 | 0.6 | Silt soil | Y | N | N |
| Nyaitho (Kenya) | (-0.179, 35.07306) | Stream | 29.1 | 7.3 | 37.9 | 0.6 | Silt soil | Y | N | N |
| Asawo (Kenya) | (-0.31817,  35.007) | River | 24.5 | 6.8 | 28.6 | 0.7 | Silt soil | Y | Y | N |
| Awach (Kenya) | (-0.23405,  34.95717) | River | 29.2 | 7.2 | 33.1 | 0.7 | Sandy soil | N | N | N |
| Awach B  (Kenya) | (-0.27203,  35.00374) | River | 29.7 | 6.4 | 38.2 | 0.6 | Sandy soil | N | N | N |
| Asao A (Kenya) | (-0.29416,  34.94199) | River | 30.1 | 7.2 | 31.2 | 0.2 | Sandy soil | N | N | N |
| Asao B (Kenya) | (-0.23403, 34.95717 | River | 31.2 | 7.5 | 33.5 | 0.3 | Sandy soil | N | N | N |
| Agoro (Kenya) | (-0.27203,  35.00374) | Stream | 30.6 | 7.5 | 32.1 | 0.2 | Silt soil | N | N | N |
| Nyando (Kenya) | (-0.17244,  34.92066) | River | 31.2 | 7.3 | 33.8 | 0.6 | Sandy soil | N | N | N |
| Ahero (Kenya) | (-0.17656,  34.92103) | Irrigation scheme | 30.6 | 7.3 | 29.1 | 1.1 | Silt soil | Y | N | N |
| Miriu (Kenya) | (-0.39753,  35.01759) | River | 31.3 | 7.6 | 32.1 | 0.5 | Sandy soil | N | N | N |
| Omondo (Kenya) | (-0.30537,  34.93082) | Stream | 30.6 | 7.3 | 39.1 | 0.2 | Silt soil | Y | N | N |
| Ochuoga (Kenya) | (-0.30075,  34.9295) | Stream | 31.3 | 7.6 | 31.2 | 0.2 | Silt soil | Y | N | N |
| Kosele (Kenya) | (-0.44471,  34.68103) | Stream | 29.8 | 7.4 | 33.6 | 1.3 | Silt soil | Y | N | N |
| Pundo (Kenya) | (-0.50641,  34.73087) | Stream | 15.3 | 6.3 | 32.5 | 1.1 | Silt soil | Y | N | N |
| Kodumo (Kenya) | (-0.41068,  34.99455) | Stream | 21.2 | 7.3 | 29.7 | 1.9 | Silt soil | Y | Y | N |
| Kodongo (Kenya) | (-0.444710, 34.681030) | Stream | 16.2 | 6.9 | 26.7 | 1.4 | Silt soil | Y | Y | N |
| Katengu (Kenya) | (-0.39756,  35.00266) | Stream | 15.2 | 7.3 | 32.1 | 1.3 | Silt soil | Y | N | N |
| K’otieno (Kenya) | (-0.51078,  34.71803) | Stream | 13.2 | 7.2 | 31.5 | 1.5 | Silt soil | Y | N | N |
| Onsando (Kenya) | (-0.71103,  35.0479) | Dam | 25.4 | 6.7 | 32.6 | 1.2 | Silt soil | Y | Y | N |
| Siwot (Kenya) | (-0.87596,  35.3683) | Dam | 15.2 | 7.2 | 32.5 | 1.1 | Sandy soil | N | N | N |
| Cheptuyet  (Kenya) | (-0.91062,  35.34843) | Stream | 25.3 | 8.3 | 27.3 | 1.1 | Silt soil | Y | Y | N |
| Kabere (Uganda) | (0.41284,  33.49619) | Stream | 19.1 | 9.2 | 27.6 | 0.4 | Silt soil | Y | N | Y |
| Kibimba (Uganda) | (0.52956,  33.85815) | Irrigation scheme | 24.7 | 7.3 | 21.7 | 1.1 | Silt soil | Y | N | N |
| Macheche (Uganda) | (0.39348,  33.48281) | Dam | 26.2 | 8.1 | 16.3 | 0.8 | Sandy soil | Y | N | Y |
| Kaitambiri (Uganda) | (1.12788,  33.70906) | Stream | 15.2 | 8.4 | 32.5 | 1.3 | Silt soil | Y | N | N |
| Nakitende (Uganda) | (1.13265,  33.67626) | Stream | 14.2 | 7.6 | 27.6 | 0.4 | Sandy soil | Y | N | N |
| Walukuba  (Uganda) | (0.44258,  33.22391) | Stream | 25.6 | 7.3 | 26.3 | 0.2 | Silt soil | Y | Y | N |
| Ariet (Uganda) | (1.16151,  33.72042) | Dam | 15.2 | 7.3 | 15.3 | 0.8 | Sandy soil | Y | N | N |
| Bisina (Uganda) | (1.59835,  33.95884) | River | 14.2 | 7.4 | 32.4 | 0.6 | Sandy soil | Y | N | N |
| Opiyai (Uganda) | (1.70238,  33.62261) | Spring | 25.2 | 6.2 | 27.2 | 0.6 | Silt soil | Y | Y | N |
| Amidakan (Uganda) | (1.93723,  33.34933) | Dam | 16.2 | 6.5 | 31.2 | 0.9 | Sandy soil | N | N | N |
| Kachung (Uganda) | (1.89908,  32.97164) | River | 15.2 | 7.4 | 38.6 | 0.5 | Sandy soil | N | N | N |
| Masindi (Uganda) | (1.69541,  32.09238) | River | 19.2 | 6.7 | 21.7 | 0.6 | Silt soil | Y | N | N |
| Kole (Uganda) | (2.302313,  32.68811) | River | 21.3 | 6.3 | 27.6 | 0.7 | Silt soil | Y | N | N |
| Masaka (Uganda) | (-0.33681,  31.71796) | Stream | 15.3 | 6.2 | 35.7 | 0.7 | Silt soil | Y | N | N |
| Kagadi (Uganda) | (0.94023,  30.81403) | Swamp | 13.2 | 8.2 | 39.8 | 0.2 | Silt soil | Y | N | N |
| Panyango (Uganda) | (2.52544,  31.46533) | River shore | 31.2 | 7.3 | 31.3 | 1.2 | Silt soil | Y | N | N |
| Muzizi (Uganda) | (0.87095,  30.72997) | River | 21.1 | 9.2 | 29.5 | 0.5 | Silt soil | Y | N | N |
| Ntoroko (Uganda) | (1.05375,  30.53696) | River | 20.8 | 8.4 | 11.2 | 0.6 | Silt soil | Y | Y | Y |
| Lubiri (Uganda) | (2.45857, 31.49975) | River | 22.1 | 7.3 | 28.3 | 0.4 | Silt soil | Y | Y | N |
| Rwizi (Uganda) | (-0.61686,  30.66828) | River | 16.2 | 7.2 | 18.6 | 0.1 | Sandy soil | Y | N | N |
| Mpanga (Uganda) | (0.658889,  30.273528) | River | 15.2 | 7.3 | 16.3 | 0.8 | Silt soil | Y | N | N |
| Kasese (Uganda) | (0.17418,  30.08111) | River | 17.2 | 7.1 | 48.6 | 0.6 | Silt soil | Y | N | N |
| Wia-Amwon (Uganda) | (2.2595,  32.87927) | Stream | 15.2 | 7.2 | 42.8 | 0.5 | Silt soil | Y | N | N |
| Wang Lei | (2.26, 32.88) | Stream | 15.2 | 7.2 | 18.6 | 0.3 | Silt soil | Y | N | Y |
| Mapea (Tanzania) | (-3.99006,  35.73814) | Irrigation scheme | 13.1 | 6.8 | 38.6 | 0.6 | Silt soil | Y | N | N |
| Njoro A (Tanzania) | (-3.32104,  37.27832) | Swamp | 16.3 | 7.3 | 11.2 | 0.6 | Silt soil | Y | N | N |
| Njoro B (Tanzania) | (-3.32598,  37.27805) | Stream | 14.3 | 6.1 | 31.7 | 0.1 | Silt soil | Y | N | N |
| Rao (Tanzania) | (-3.3381,  37.35776) | River | 15.2 | 8.5 | 32.1 | 0.5 | Silt soil | Y | N | N |
| Kikavu (Tanzania) | (-3.43637, 37.29482) | Irrigation scheme | 16.2 | 8.2 | 12.4 | 0.2 | Sandy soil | Y | N | Y |
| Mabogini (Tanzania) | (-3.40061,  37.36256) | Irrigation scheme | 24.3 | 8.8 | 27.6 | 0.7 | Silt soil | Y | N | N |
| Chiwe (Tanzania) | (-6.21983,  36.74357) | River | 15.2 | 7.3 | 29.6 | 0.3 | Sandy soil | Y | N | N |
| Ihanda (Tanzania) | (-6.24869,  36.71776) | Stream | 15.2 | 7.2 | 32.7 | 0.2 | Silt soil | Y | N | N |
| Mhango (Tanzania) | (-6.24053,  37.5274) | Stream | 14.2 | 8.5 | 35.2 | 0.3 | Silt soil | Y | N | N |
| Mvomero (Tanzania) | (-6.30143,  37.44623) | River | 16.2 | 8.2 | 31.6 | 0.5 | Sandy soil | Y | N | N |
| Mkindu (Tanzania) | (-6.24803,  37.54699) | River | 14.2 | 7.5 | 33.7 | 0.6 | Sandy soil | N | N | N |
| Mkindo (Tanzania) | (-6.26084,  37.55375) | Irrigation scheme | 15.2 | 7.2 | 12.4 | 1.1 | Silt soil | Y | N | N |
| Mlegeni (Tanzania) | (-6.89333,  37.07764) | Stream | 16.3 | 7.6 | 31.2 | 0.7 | Silt soil | Y | N | N |
| Dago (Tanzania) | (-6.8934,  37.07775) | Stream | 15.3 | 8.1 | 32.1 | 0.4 | Silt soil | Y | N | N |
| Zombo (Tanzania) | (-6.96513,  36.91566) | Irrigation scheme | 16.2 | 8.5 | 36.6 | 0.6 | Silt soil | Y | N | N |
| Tindiga (Tanzania) | (-6.89507,  37.0883) | Irrigation scheme | 16.2 | 8.2 | 37.6 | 0.7 | Silt soil | Y | N | N |
| Chabi (Tanzania) | (-7.31147,  36.52885) | Irrigation scheme | 16.3 | 8.7 | 36.6 | 0.2 | Silt soil | Y | N | N |
| Mwega (Tanzania) | (-7.31152,  36.52878) | Irrigation scheme | 13.2 | 8.1 | 37.4 | 0.9 | Silt soil | Y | Y | N |
| Bwawani (Tanzania) | (-7.73513,  35.71792) | Dam | 24.1 | 8.5 | 18.6 | 0.7 | Silt soil | Y | N | N |
| Ilongo (Tanzania) | (-8.76628,  33.74477) | Irrigation scheme | 17.2 | 7.1 | 32.7 | 0.3 | Silt soil | Y | N | N |
| Nsalagah (Tanzania) | (-8.88944,  33.57027) | Stream | 16.3 | 7.6 | 37.5 | 0.3 | Silt soil | Y | N | N |
| Migombani (Tanzania) | (-9.31781,  32.76955) | Stream | 32.2 | 7.3 | 34.6 | 0.1 | Silt soil | Y | N | N |
| Kasekera (Tanzania) | (-4.689278,  29.622083) | Stream | 31.2 | 7.2 | 39.2 | 0.5 | Silt soil | Y | N | N |
| Singida (Tanzania) | (-4.788596, 34.745608) | River | 31.2 | 7.5 | 33.2 | 0.5 | Sandy soil | N | Y | N |
| Kibilizi (Tanzania) | (-4.860920, 29.624456) | Stream | 26.1 | 7.3 | 34.3 | 0.3 | Silt soil | N | N | N |
| Musoma (Tanzania) | (-1.494590, 33.810970) | River | 29.2 | 6.5 | 39.5 | 0.3 | Sandy soil | Y | N | N |
| Bweri (Tanzania) | (-1.537530, 33.854760) | Stream | 14.7 | 6.9 | 33.2 | 0.4 | Silt soil | Y | N | N |
| Ziro ziro (Tanzania) | (-1.598370, 33.909480) | Stream | 31.2 | 7.2 | 35.4 | 0.3 | Silt soil | Y | N | N |
| Musoma (Tanzania) | (-1.414650, 34.213500) | Stream | 29.2 | 7.5 | 33.6 | 0.6 | Sandy soil | Y | N | N |
| Kongoro (Tanzania) | (-1.339650, 34.385780) | Stream | 15.8 | 6.9 | 34.2 | 0.4 | Sandy soil | Y | N | N |
| Suguti (Tanzania) | (-1.680613, 33.703669) | River | 16.5 | 7.1 | 35.2 | 0.2 | Sandy soil | Y | N | N |
| Nansimo (Tanzania) | (-2.157424, 33.443241) | River | 16.3 | 7.9 | 32.6 | 0.5 | Sandy soil | Y | N | N |
| Lamadi (Tanzania) | (-2.246043, 33.836968) | River | 15.3 | 7.2 | 38.6 | 0.4 | Sandy soil | Y | N | N |
| Mwamanyili (Tanzania) | (-2.372647, 33.563318) | River | 14.3 | 7.1 | 35.3 | 0.9 | Silt soil | Y | N | N |
| Igogo (Tanzania) | (-2.539848, 32.901068) | River | 29.3 | 6.8 | 35.2 | 0.8 | Silt soil | Y | N | N |
| Mnyuzi (Tanzania) | (-5.305361, 38.628585) | Stream | 31.2 | 6.5 | 32.5 | 0.6 | Silt soil | Y | Y | N |
| Utwigu (Tanzania) | (-4.446061, 33.050964) | Stream | 20.2 | 6.3 | 29.6 | 0.6 | Silt soil | Y | N | N |
| Chigunga (Tanzania) | (-2.778439, 32.034190) | River | 14.2 | 6.5 | 37.3 | 0.7 | Sandy soil | Y | N | N |
| Igombe (Tanzania) | (-4.888247, 32.754505) | Dam | 28.3 | 7.2 | 38.5 | 2.6 | Sandy soil | N | N | N |
| Mwaseni (Tanzania) | (-7.792478, 38.094281) | River | 17.2 | 7.5 | 32.1 | 0.6 | Sandy soil | Y | N | N |
| Songea (Tanzania) | (-10.662715, 35.56577) | Dam | 15.5 | 6.9 | 37.3 | 0.8 | Sandy soil | Y | N | N |
| Masonya (Tanzania) | (-10.6992, 37.356541) | River | 15.4 | 8.1 | 32.5 | 0.4 | Sandy soil | Y | N | N |
| Shanwe (Tanzania) | (-6.32568, 31.052803) | Stream | 15.2 | 7.3 | 37.5 | 0.7 | Silt soil | Y | N | N |
| Sangari (Tanzania) | (-5.17815, 31.15624) | River | 16.3 | 6.5 | 35.4 | 0.8 | Silt soil | N | N | N |
| Ntalikwa (Tanzania) | (-5.07492, 32.71561) | Stream | 17.3 | 7.4 | 32.1 | 0.3 | Sandy soil | Y | N | N |
| Ntalikwa (Tanzania) | (-5.0742, 32.71541) | Irrigation scheme | 16.7 | 7.1 | 39.1 | 0.6 | Silt soil | Y | N | N |
| Sumbawanga (Tanzania) | (-7.99843, 31.61841) | Stream | 15.6 | 6.7 | 33.2 | 0.4 | Silt soil | Y | N | N |
| Bujonde (Tanzania) | (-9.66072,  33.95171) | River | 30.4 | 7.2 | 38.1 | 0.6 | Sandy soil | Y | N | N |
| Matema (Tanzania) | (-9.49525,  34.02471) | River | 29.3 | 6.9 | 34.2 | 0.4 | Sandy soil | Y | N | N |
| Igalula (Tanzania) | (-5.63833,  32.62784) | Stream | 28.1 | 7.4 | 37.2 | 0.5 | Silt soil | Y | Y | N |
| Mwamgongo (Tanzania) | (-4.62742,  29.65162) | Stream | 25.9 | 6.8 | 28.1 | 0..7 | Silt soil | Y | N | N |
| Bugamba (Tanzania) | (-4.56683,  29.65203) | Stream | 32.1 | 8.1 | 35.7 | 0.2 | Silt soil | Y | N | N |
| Kiziba (Tanzania) | (-4.51933, 29.66061) | Stream | 29.1 | 7.2 | 36.2 | 0.6 | Silt soil | Y | N | N |
| Babati (Tanzania) | (-4.227588, 35.744211) | River | 24.5 | 8.1 | 53.2 | 0.2 | Silt soil | Y | Y | N |
| Mbaka (Tanzania) | (-9.549804, 33.957001) | River | 29.5 | 6.8 | 35.2 | 0.9 | Silt soil | Y | N | N |
| Mwaya (Tanzania) | (-9.558547, 33.948174) | Stream | 31.4 | 7.2 | 32.1 | 0.6 | Silt soil | Y | N | N |
| Galana (Tanzania) | (-2.201967, 38.058157) | River | 32.3 | 7.3 | 33.2 | 0.5 | Silt soil | Y | N | N |
| Mpanda (Tanzania) | (-6.355559, 31.053331) | River | 29.3 | 7.1 | 35.3 | 0.4 | Silt soil | Y | N | N |
| Ibulwa (Tanzania) | (-5.215938, 32.669059) | River | 31.4 | 6.8 | 36.2 | 0.9 | Silt soil | Y | N | N |
| Shama (Tanzania) | (-6.485610, 33.110264) | River | 15.3 | 7.3 | 36.2 | 0.4 | Silt soil | Y | N | N |
| Lupa (Tanzania) | (-7.904587, 33.303941) | River | 13.1 | 6.7 | 32.2 | 0.6 | Sandy soil | Y | N | N |
| Mindu (Tanzania) | (-6.868224, 37.613939) | Dam | 29.2 | 7.2 | 33.2 | 0.6 | Sandy soil | Y | N | N |
| Midiho (Tanzania) | (-5.538566, 32.590210) | River | 17.3 | 7.5 | 35.2 | 0.6 | Sandy soil | Y | N | N |
| Kisisi (Tanzania) | (-5.244343, 32.707229) | River | 15.3 | 7.9 | 33.2 | 0.5 | Sandy soil | Y | N | N |
| Duluti (Tanzania) | (**-**3.385123, 36.786488) | Dam | 13.2 | 9.1 | 39.6 | 0.4 | Sandy soil | Y | N | N |
| Holili (Tanzania) | (-3.378823, 37.620629) | Stream | 17.3 | 8.3 | 39.2 | 0.2 | Silt soil | Y | N | N |
| Mkondoa (Tanzania) | (-6.825841, 37.161774) | River | 15.3 | 6.3 | 32.3 | 0.8 | Sandy soil | Y | N | N |
| Mkundi (Tanzania) | (-6.384851, 37.360174) | River | 17.6 | 7.1 | 34.2 | 0.9 | Silt soil | Y | N | N |
| Gairo (Tanzania) | (-6.127093, 36.867884) | Stream | 29.7 | 7.2 | 38.1 | 0.4 | Silt soil | Y | N | N |
| Lumuma (Tanzania) | (-7.333937, 36.521680) | River | 29.5 | 7.3 | 35.1 | 0.4 | Sandy soil | Y | N | N |
| Bomani (Tanzania) | (-1.345125, 34.372426) | Stream | 31.2 | 7.1 | 38.2 | 0.7 | Silt soil | Y | N | N |
| Mori (Tanzania) | (-1.346410, 34.087468) | River | 30.2 | 6.4 | 37.2 | 0.3 | Sandy soil | Y | N | N |
| Nkumbu (Tanzania) | (-3.695077, 33.460108) | River | 31.2 | 7.9 | 35.1 | 0.9 | Silt soil | Y | N | N |
| Mori (Tanzania) | (-3.695077, 33.460108) | River | 29.4 | 8.1 | 35.2 | 0.4 | Silt soil | Y | N | N |
| Nkaiti (Tanzania) | (-3.791196,  35.9164310) | River | 31.2 | 9.2 | 33.1 | 0.3 | Sandy soil | Y | N | N |
| Pangani (Tanzania) | (-3.382294, 37.324104) | River | 30.8 | 7.4 | 35.2 | 0.3 | Sandy soil | Y | N | N |
| Nyankanga (Tanzania) | (-1.588600, 33.901086) | Stream | 29.2 | 7.2 | 36.2 | 0.8 | Sandy soil | Y | N | N |
